# A Comparison of Persistence of SARS-CoV-2 Variants on Stainless Steel

**DOI:** 10.1101/2021.04.08.438833

**Authors:** Thomas Pottage, Isobel Garratt, Okechukwu Onianwa, Antony Spencer, Susan Paton, Jake Dunning, Allan Bennett

## Abstract

The survival of newer variants of SARS-CoV-2 on a representative surface has been compared to the established UK circulating isolate to determine whether enhanced environmental stability could play a part in their increased transmissibility. Stainless-steel coupons were inoculated with liquid cultures of the three variants, with coupons recovered over seven days and processed for recoverable viable virus using plaque assay. After drying, there was no significant difference in inactivation rates between variants. Indicating there is no increased environmental persistence from the new variants.

## Introduction

On 11^th^ March 2020, the World Health Organization declared the recently emerged SARS-CoV-2 virus had resulted in a pandemic of COVID-19. Since that time, waves of COVID-19 activity have been seen globally and continue to occur one year later. In late 2020 two SARS-CoV-2 variants of concern emerged, the so-called Kent variant (lineage B.1.1.7 or VOC202012/01) and a South African variant (lineage B.1.351 or 20H/501Y.V2), both of which have been detected in multiple countries, including the UK [1]. Both variants have the N501Y mutation, which, along with other mutations affecting the receptor binding domain of the spike glycoprotein, are associated with increased host receptor-binding affinity; this may, in part, result in increased transmissibility [2–4].

Surfaces can potentially be contaminated with SARS-CoV-2 directly from the deposition of particles released from an infected person’s respiratory tract or from contaminated hands, potentially leading to transfer through contact. Agents remaining viable for longer are more likely to be transferred and cause infection. Environmental sampling has detected SARS-CoV-2 RNA on environmental surfaces in healthcare spaces and this evidence, coupled with laboratory studies demonstrating viral viability on surfaces, has contributed to the assumption that environmental contamination and fomite spread play a role in viral transmission [5, 6].

It is important to understand if the increase in transmissibility observed with the new variants is solely from spike mutations leading to increased host receptor affinity, or if the mutations have an impact on environmental persistence. These factors could have implications for infection control to limit onwards transmission of the virus. This study was designed to determine possible differences in surface persistence between the previously established circulating strain of SARS-CoV-2 hCOV-19/England/2/2020 (EPI_ISL_407073 England) and the new emergent strains B.1.1.7 and B.1.351. This study utilised stainless steel coupons and plaque assays to investigate the viability of these SARS-CoV-2 isolates.

## Materials and methods

### SARS-CoV-2 isolates

Three SARS-CoV-2 variants were isolated and propagated for use in this study. The variant England 02/2020 HCM/V/052 (EPI_ISL_407073 England) was prepared as described by Paton *et al* [6].

Human nCoV19 isolate/England/MIG457/2020 (lineage B.1.1.7 or VOC202012/01) was isolated by the Public Health England Medical Interventions Group (MIG) in hSLAM cells. Human nCoV19 isolate/England/H204661641/2020 (lineage B.1.351 or 20H/501Y.V2) was isolated in Vero-E6 cells at the Barclay Laboratory, Imperial College London, and transferred to PHE. After isolation, both variants were passaged twice further in hSLAM cells then centrifuged at 1,000 g for 10 minutes to produce stocks at a concentration of 3.1 × 10^6^ pfu/ml (B.1.1.7) and 1.7 × 10^6^ pfu/ml (B.3.5.1). All work handling SARS-CoV-2 was performed within a containment level 3 laboratory.

### Experimental procedure

Coupon preparation, inoculation, exposure and recovery was completed as documented previously [6]. Briefly, triplicate stainless-steel coupons for each time point and variant were inoculated with 10 μl (×2) droplets of the variant stock suspension within a flexible film isolator (FFI). Triplicate samples per variant were taken by placing coupons individually, into 1 ml of complete minimal essential medium (cMEM) with four glass beads (3mm diameter). Seven time points used were; 0 time point (immediately after inoculation), 2.5 hours (once completely dried), and subsequent time points (24, 48, 72, 96 and 168 hours), coupons were kept within the FFI for their exposure period. During the study the average temperature was 19 °C and relative humidity was 57 %. Samples were vortex mixed for one minute before freezing at −80 °C. The recovery media was thawed to room temperature, serially diluted and plaque assayed on Vero-E6 cells, before staining and enumeration of plaques.

### Statistical Analysis

All statistical analyse and data manipulation were completed in STATA (v16.1).

## Results and discussion

During the initial drying period of the inoculum on stainless-steel coupons all three isolates exhibited an initial sharp drop in viability, which subsequently further decreased after the 2.5 hour drying period. Of the initial reductions, the common predecessor (EPI_ISL_407073) reduced by approximately one log_10_, whilst a reduction of 0.15log_10_ and <0.1log_10_ for the B.1.1.7 and B.1.351 variants were seen (Figure 1). The rate of decrease in viability reduced after the drying process, with all variants following a similar trend in recovery of viable virus. However, during the drying process, there was a significant difference between the units recovered of common predecessor virus and the B.1.1.7 and B.1.351 variants (*p*=0.01), whereas there was no significant difference between the units recovered of the B.1.1.7and B.1.351 variants (*p*=0.25).

**Figure 1.**
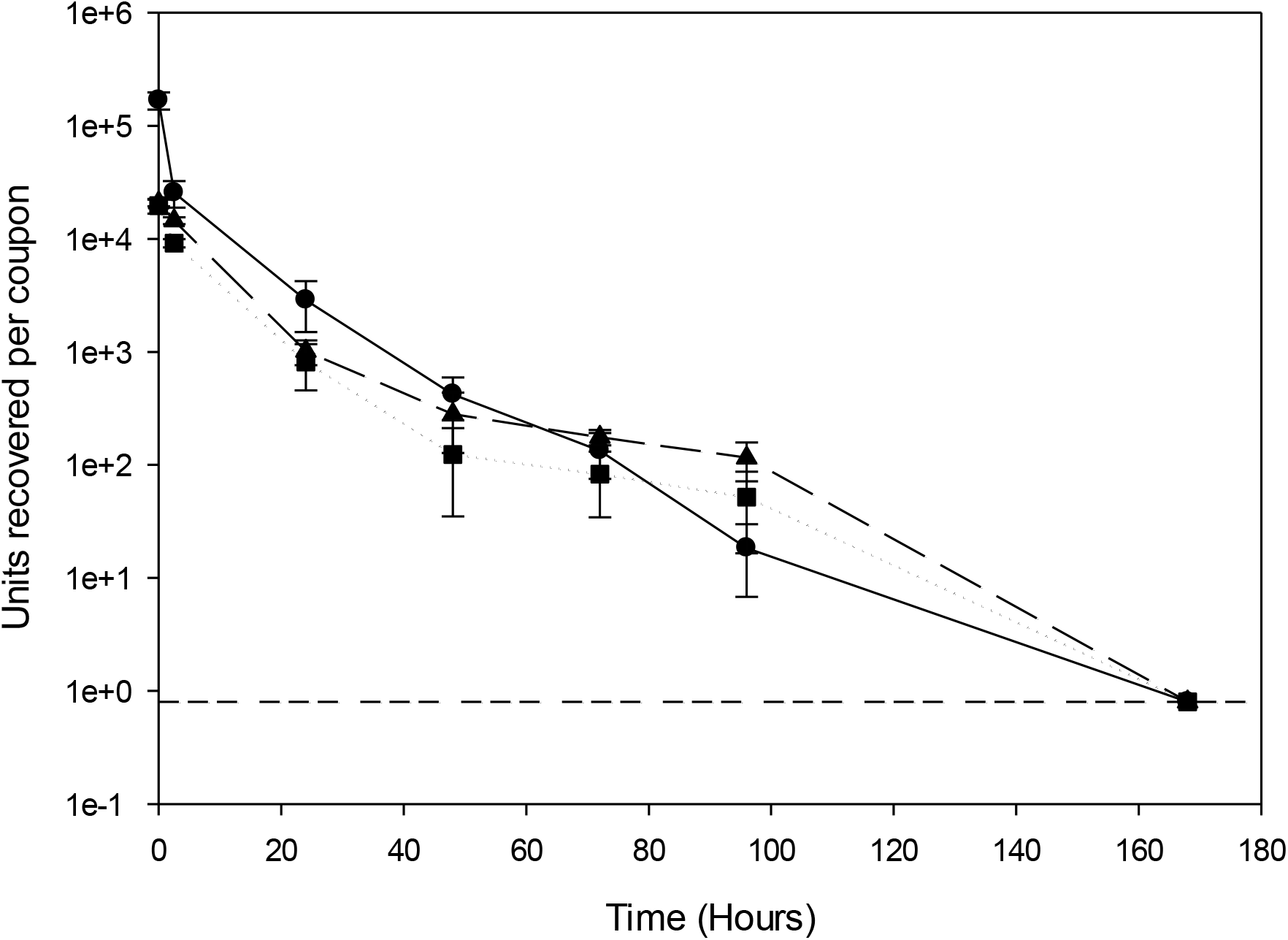
Mean quantities of viable virus recovered (pfu/coupon) from stainless-steel loaded with England 02/2020 HCM/V/052 (circles), Kent variant (triangles) and South African variant (squares). Error bars represent the standard deviation from three replicates. Dashed line represents the limit of detection of the plaque assay for the combined assays from the triplicate coupons.

After completion of the drying process (2.5 hours), the rate in reduction in viability of the three variants was not significantly different from each other (*p*=0.12). Indicating, once dried, the variants lose viability at a similar rate, with no viable virus detected after 7 days. This similarity in viability of the variants over time shows that the likelihood of a contaminated surface being touched and viable virus being present is the same for the three variants studied.

To the best of our knowledge, this is the first study comparing persistence of SARS-CoV-2 variants on a representative environmental surface in an experimental setting. Previous studies have investigated differences between SARS-CoV-2 and other coronaviruses. Van Doremalen *et al* reported the persistence of SARS-CoV-2 and SARS-CoV-1 was similar on stainless-steel, observing a >3 log reduction in viability within 48 hours, from the starting inoculum reducing to the detection limit in this time [5]; suggesting that these results showed contaminated fomites were a potential transmission route. Sizun *et al*. compared two human coronaviruses (229E and OC43), with 229E surviving longer than OC43 after drying [7]. No indication was given to explain the differences in the persistence between the coronaviruses. Other studies have focused on single variants/isolates, and then compared data obtained with those from other groups. The challenges to this approach are the variations in methods between studies, making comparison difficult [8, 9]. Incubating dried SARS-CoV-2 virus at a higher temperature than room temperature (28°C versus 20-25°C) increased the inactivation rate, but viable virus was recovered over the same time period at both temperatures [10].

This study used high concentration suspensions of SARS-CoV-2 variants to allow that the detection of small differences in recovered viability. These starting concentration levels have not been found during environmental surface sampling, so are artificially high in comparison, but using a lower starting concentration could preclude the detection of inter-strain differences through the rapid inactivation of the virus on the surface, giving few data points for analysis. Stainless-steel was chosen as the carrier material for this study because it is representative of touchpoints that will be frequently contacted by different individuals and provides an inanimate surface on which SARS-CoV-2 has been shown to persist. This is in comparison to other surfaces such as face masks [6, 8], which are likely to be contacted by fewer individuals and disposed of after use. Given that multiple environmental factors can influence virus SARS-CoV-2 viability [9, 10], the persistence of viable virus on surfaces and the associated transmission risk may be considerably shorter in various real-life settings.

In conclusion, the results from this study show that during the initial drying period there is a greater reduction in the viability of the common predecessor virus compared to the B.1.1.7 and B.1.351 variants, but once dried, the rates of inactivation are very similar. Consistent with previous findings [6], viable virus is not recoverable from stainless-steel coupons after seven days, under experimental conditions.

## Acknowledgements

Dr K Bewley in Medical Interventions Group, PHE, Porton Down, for producing the viral Human nCoV19 isolate/England/MIG457/2020 (lineage B.1.1.7 or VOC202012/01) and Human nCoV19 isolate/England/H204661641/2020 (lineage B.1.351 or 20H/501Y.V2) stocks for this study.

Professor W Barclay, Barclay Laboratory, Imperial College London, for providing the Human nCoV19 isolate/England/H204661641/2020 (lineage B.1.351 or 20H/501Y.V2) variant isolate.

Dr K Richards in High Containment Microbiology, PHE, Porton Down, for producing the England 02/2020 HCM/V/052 (EPI_ISL_407073 England) viral stock for this study.

The views expressed in this article are those of the authors and are not necessarily those of PHE or the Department of Health and Social Care. This manuscript is Crown copyright and is reproduced with the permission of Public Health England and the Controller of HMSO.

